# Chromatin structure shapes the search process of transcription factors

**DOI:** 10.1101/050146

**Authors:** Neslihan Avcu, Nacho Molina

## Abstract

The diffusion of regulatory proteins within the nucleus plays a crucial role in the dynamics of transcriptional regulation. The standard model assumes a 3D plus 1D diffusion process: regulatory proteins either move freely in solution or slide on DNA. This model however does not considered the 3D structure of chromatin. Here we proposed a multi-scale stochastic model that integrates, for the first time, high-resolution information on chromatin structure as well as DNA-protein interactions. The dynamics of transcription factors was modeled as a slide plus jump diffusion process on a chromatin network based on pair-wise contact maps obtained from high-resolution Hi-C experiments. Our model allowed us to uncover the effects of chromatin structure on transcription factor occupancy profiles and target search times. Finally, we showed that binding sites clustered on few topological associated domains leading to a higher local concentration of transcription factors which could reflect an optimal strategy to efficiently use limited transcriptional resources.

## Introduction

Gene expression is regulated by transcription factors (TFs) that recognize specific regulatory DNA sequences [1]. The search strategies that TFs use to find these regulatory target sites are key to understand the dynamics of transcriptional regulation. The standard model assumes a 3D plus 1D diffusion process: TFs either move freely in solution or slide on DNA [2, 3, 4]. However recent single moleculeexperiments have suggested that 3D structure of chromatin influences the diffusion process [5]. Here we proposed a new multi-scale stochastic model that integrates, for the first time, high-resolution information on the 3D structure of chromatin as well as DNA-protein interactions. This model allowed us to provide genome-wide testable predictions for TF occupancy and target search time of any locus in the genome.

**Figure 1:**
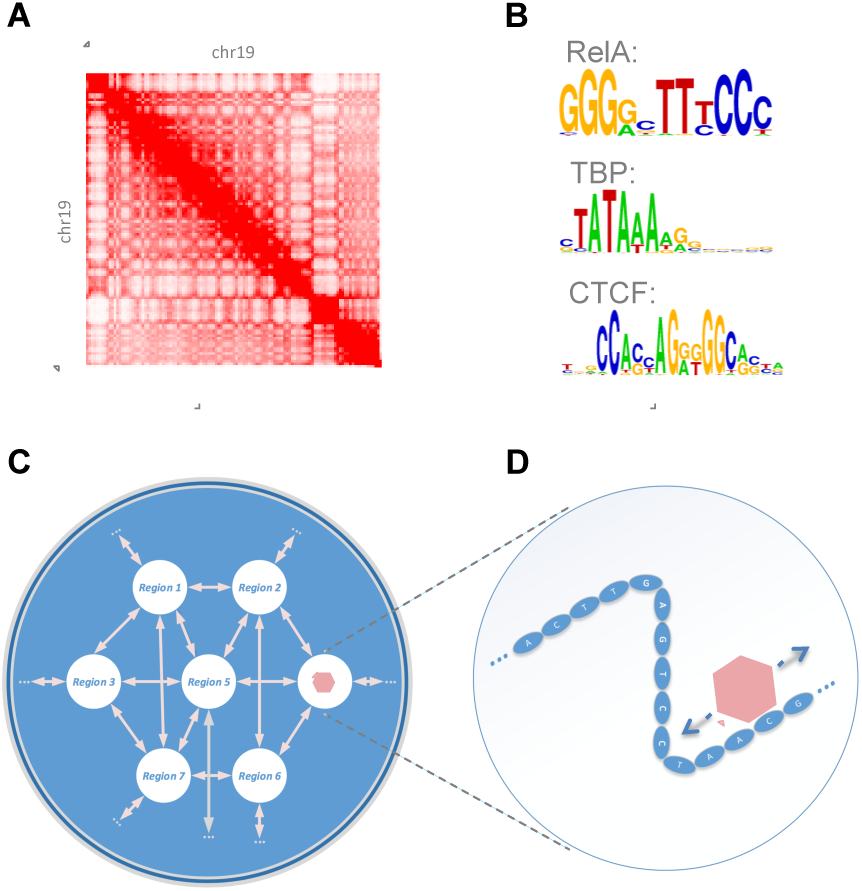
Model of TF diffusion integrating chromatin structure information. (A) Inter-chromosome interactions produced by *in-situ* Hi-C method [6]. Intensity corresponds to normalized number of reads. (B) Sequence motifs representing DNA-protein affinity for three TFs [7]. (C) Diagram of the TF diffusion multi-scale model. Chromatin is represented as a network where nodes are genomic regions (5kb) and edges are pair-wise chromatin interactions derived from Hi-C contact maps. TF can jump from one genomic region to another if there is an edge that connects both regions or slide on DNA taking into account specific DNA sequence interactions according to the TF motif.

## TF diffusion dynamics as a random walk on a chromatin network

To model the dynamics of a TF within a chromatin structure we proposed a jump and slide diffusion model. We assumed that the diffusive protein can slide on the DNA of a given genomic region or alternatively jump into a new region if both are close in space. Hi-C method [8, 9, 6] provides precisely the information of which genomic regions are likely to be physically in contact. Based on recently published high-resolution contact maps of the mouse genome [6], we built a network where nodes represent genomic regions of 5kb and edges are pair-wise chromatin interactions. The diffusion process is then described as a random walk on the resulting chromatin network where the diffusive protein stays in each region a characteristic residence time τ_i_ before jumping into a new one. We assumed that the residence times τ_i_ are the result of a sliding process at the base-pair resolution within the 5kb regions where specific and non-specific DNA-interactions were model based on available position wight matrices (PWMs) [7] (see supporting material).

Note that the key parameters of our diffusion model are therefore the topology of the network, which describes the chromatin structure, and the residence times τ_i_ that encodes the sequence-specific interactions. We believe this is the simplest genome-wide model that is able, for the first time, to incorporate all the information we have on both the chromatin structure and on specific DNA-protein interactions.

## TF occupancy is proportional to residence time and contact degree

The time-evolution of the probability of finding the diffusive protein at a certain genomic region can be described by a master equation (see supporting material). Interestingly, although the network topology representing the chromatin structure can be very complex the steady-state solution can be solved analytically leading to a very simple expression. The occupancy of a region *i* is proportional to the residence time τ_*i*_ and the contact degree *d*_*i*_, namely the number of physical contacts with other regions. That is,

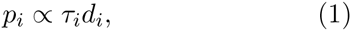

where the proportionality factor is just the normalization constant. This nicely shows that the occupancy of a region is the result of two independent factors: the first summarizes the DNA-protein interaction and the sliding process (τ_i_); and, the second describes the effects of the chromatin structure (*di*). Remarkably, only a local structure property, namely the number of contacts of a given region, plays a role in determining its occupancy.

To test our model we compared the predicted genome-wide occupancy profiles of three different transcription factors (RelA, TBP and CTCF) with available ChIP-seq data [10]. As shown in Fig. 2 (left panels), TF occupancy strongly correlates with contact degree. Indeed, contact degrees span for one or two orders of magnitude causing a similar span in occupancy levels. Interestingly, genomic regions with TF binding sites (TFBSs) show a significantly large number of contacts compared regions without sites. Indeed, we observed an increase of 38%, 50%and 33% in the average contact degree of regions with binding sites for RelA, TBP and CTCF respectively.

**Figure 2:**
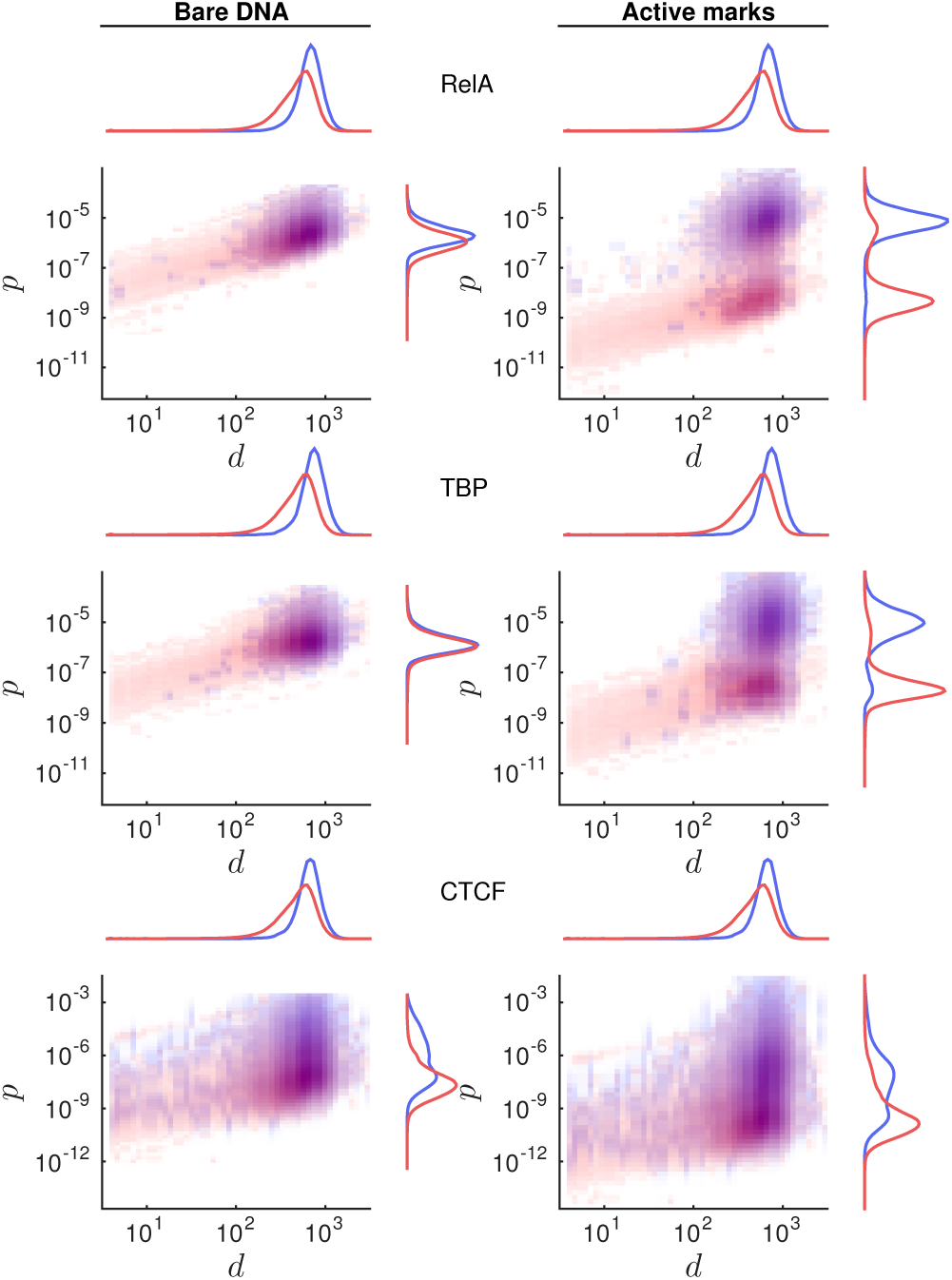
TF occupancy is proportional to contact degree and residence time. 2D histograms showing fraction of genomic regions with certain TF occupancy and contact degree. Blue and red histograms were obtained using genomic regions with and without TFBSs measured by ChIP-seq. Left panels show TF occupancies assuming bare DNA and right panels TF occupancies when active histone marks are used to modulate residence times. 1D histograms after marginalization are shown on top and right side of each panel. Regions with TFBSs show a higher occupancy and contact degree than regions without TFBSs. Active histone marks cause a drastic increase in the expected occupancy level of regions with TFBSs (*h*= 10-^3^).

Although the structural factor in Eq. 1 plays a significant role on determining the occupancy levels comparable to the sequence factor, it does not fully discriminate regions with and without TFBSs (Fig. 2). To improve the predicted occupancy we recalculated the residence times τ_*i*_ taking into account profiles of active histone marks (H3K4me1, H3K4me3 and H3K27Ac) measured using ChIP-seq [10]. We assume that DNA-protein interactions are reduced by a factor *h* on DNA that shows low levels of the three active marks. The new predicted occupancy for RelA and TBP clearly discriminate between regions with and without binding sites for RelA and TBP (Fig. 2, right panels).

## Chromatin structure reduces TF search time

Compared to previous thermodynamic approaches [12], our stochastic model allowed us to go beyond the steady-state solution. Indeed, we were able to calculate the expected search time *t*_*s*_ required for a TF to find any given genomic region. In this context, the search process is a random walk on the chromatin network until the TF hits for the first time the region of interest. The search time then follows a phase-type distribution whose mean is related to the inverse of the transition rate matrix of the random walk and the probability distribution over regions where the search could be initiated (see supporting material). First, it is easy to show that *t*_*s*_ is proportional to the average residence time (τ).Intuitively, the stickier the TF-DNA interaction is the longer the time is required to find aparticular locus. Note that average residence times have been measured by single-molecule mi-croscopyshowing values for different TFs in the range of seconds [13, 11].

The matrix inversion required to calculate *t*_*s*_ can be computationally very costly especially for large matrices. Therefore we first considered a search process only on chromosome 19, the smallest one. The times required for RelA to find genomic regions of chromosome 19 are shown in Fig 3A. Strikingly, search times are inversely proportional to the contact degree *d* of the searched region. Therefore, regions with RelA binding sites are found faster as they show larger contact degrees (blue dots in Fig. 3A). Notice that the search time predicted by the 3D plus 1D diffusion model is the same for all regions in the genome and, interestingly, larger than the one obtained with our model for most of the regions with TFBSs (dashed line in Fig. 3A and supporting material). These results show that chromatin structure play a crucial role in shaping the search process of transcription factors.

**Figure 3:**
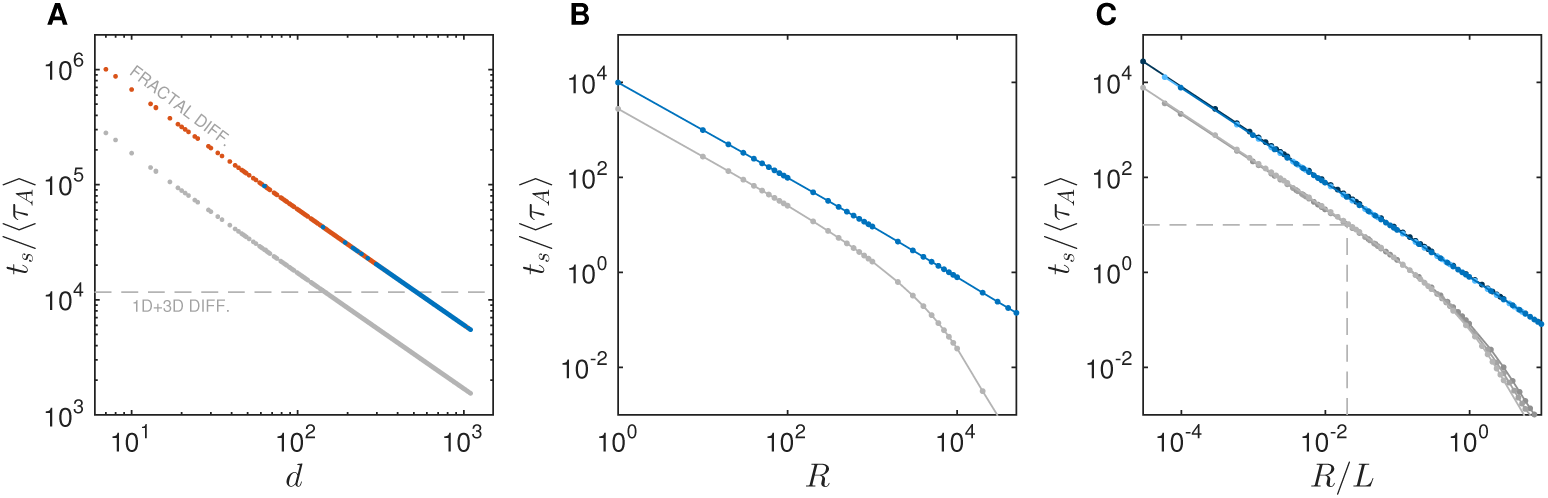
Target search time is inversely proportional to contact degree. (A) Relationship between contact degree *d* and search time *t*_*s*_, normalized by the average residence time on active chromatin❬τ_*A*_❭, of genomic regions in chromosome 19 with (blue) and without (red) TFBSs (no histone marks were considered and therefore ❬τ_*A*_❭=❬τ_*I*_❭). In grey predicted search timeswhen active histone marksareconsidered and ❬τ_*A*_❭/❬τ_*I*_❭≈10^3^ leading to a 28% reduction. (B) Relationship between number of TF molecules*R* and the search time of a given genomic region in chromosome 19 assuming bare DNA (blue) or active-marks modulation (grey). (C) Same as in B but considering the whole genome at different cross-grained scales assuming bare DNA (dark blue: 75kb; blue: 100kb; light blue: 200kb) or active-marks modulation (grey blue: 75kb; grey: 100kb; light grey: 200kb). As an example: *R*= 10^4^ molecules searching within a chromatin network of *L* = 5. 10^5^ genomic regions of length 5kb find the target region in 1s if we assumed ❬τ_*A*_❭ =10s [11] (gray dashed lines).

Next, we investigated how active histone marks affected the search process. Notice that the average residence time can be expressed as a weighted sum of the average residence times in active and inactive regions: ❬τ❭ = ❬ τ_*A*_❭*ƒ*_*A*_ + ❬τ_*I*_❭*ƒ*_*I*_, where the weights *ƒ*_*A*_ and *ƒ*_*I*_ are the fraction of times that the TF visits active and inactive regions respectively (see supporting material). Assuming that the affinity of the TF-DNA interactions on active regions was three orders of magnitude larger than on inactive region (❬τ_*A*_❭/❬τ_*I*_❭ ⋍ 10^3^) [14, 15], the search time was reduced by 28% (see Fig 3A) as ❬τ❭ ⋍ ❬τ_*A*_❭*ƒ*_*A*_. This speed up of the search process iscaused by an effective chromosome length reduction due to the low affinity of non-active regions.

Up to now we considered the process of a single TF searching for a given region. A more relevant biological scenario is to consider *R* molecules that perform the search independently. In this case, it is easy to show that the resulting search time is the probability that *R* - 1 molecules did not find the region after certain time *t*_*s*_ and one molecule did find it at precisely *t*_*s*_ (see supporting material). The relationship between the search time *t*_*s*_ and the number of molecules *R* is shown in Fig. 3B for a region of the chromosome 19 that contains RelA sites. Consistently with what one would obtained from reaction kinetics based on the law of mass action, *t*_*s*_ is inversely proportional to the number of molecules *R*. Strikingly, when we took into account active mark profiles we found a regime for large number of molecules (*R* > 10^3^) at which the search time decreases faster with the number of searching molecules. This results indicates that the combination of chromatin structure and histone marks increases the efficiency of the search process. This has an important consequence on how the kinetics of gene activation depends on the concentration of the regulatory proteins.

To extend our model to consider the whole genome in a tractable manner we coarse-grained the chromatin network at different resolutions to effectively reduce its size. Remarkably, different resolutions (200kb, 100kb and 75kb) lead to comparable results when we considered the concentration of molecules respect to the chromatin network size *L* instead of absolute molecules numbers (see Fig. 3C). This allowed us to extrapolate our results and obtain search times considering the whole genome at high resolution. For instance, the genomic region considered in Fig. 3B is found on average in 10 seconds when 10^4^ molecules are searching in parallel in the whole chromatin network with approximately 5. 10^5^ genomic regions of 5kb length.

The previous results, all together, lead to the following expression for the search time:

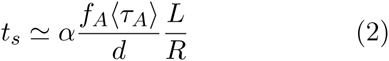

where the structure constant *α* encodes global properties of the chromatin network topology, however does not depend on the searched region (see supporting material). Notice that when a promoter region is considered, the inverse of the search time can be interpreted as an effective ‘on’ rate: *k*_on_ = 1/t_s_, which incorporates the influence of the chromatin structure on TF binding rates.

## Asymmetric distribution of TFBSs among TADs traps TF diffusion

Large local chromatin interaction domains, termed topological associated domains (TADs) have been identified across cell types and organism [9]. Next we studied the role of these higher order chromatin structures on the diffusion dynamics. In order to do that, first, we computed the distribution of binding sites across TADs of 25 TFs involved in the immune response [10]. Remarkably, Fig 4A shows that the number of TFBSs is strongly over-represented in few TADs compared with a random distribution. Furthermore, histograms showing the frequency of TADs with certain number of TFBSs show that domains both with few or many bindings sites occur more often than expected by a random model (See Fig. 4B-C and Fig. S[#]). These results suggest that the concentrations of TFs is not homogenous across the nucleus as TADs with large number of binding sites trap TFs for longer times. To test this we calculated using our diffusion model the escaping time *t*_e_ of RelA from 2970 TADs previously identified [6]. As expected domains with many binding sites show 3 fold increase in the escaping time on average (Fig. 4D). This difference disappeared when we randomized the residence times τ_i_ (Fig. 4E).

**Figure 4:**
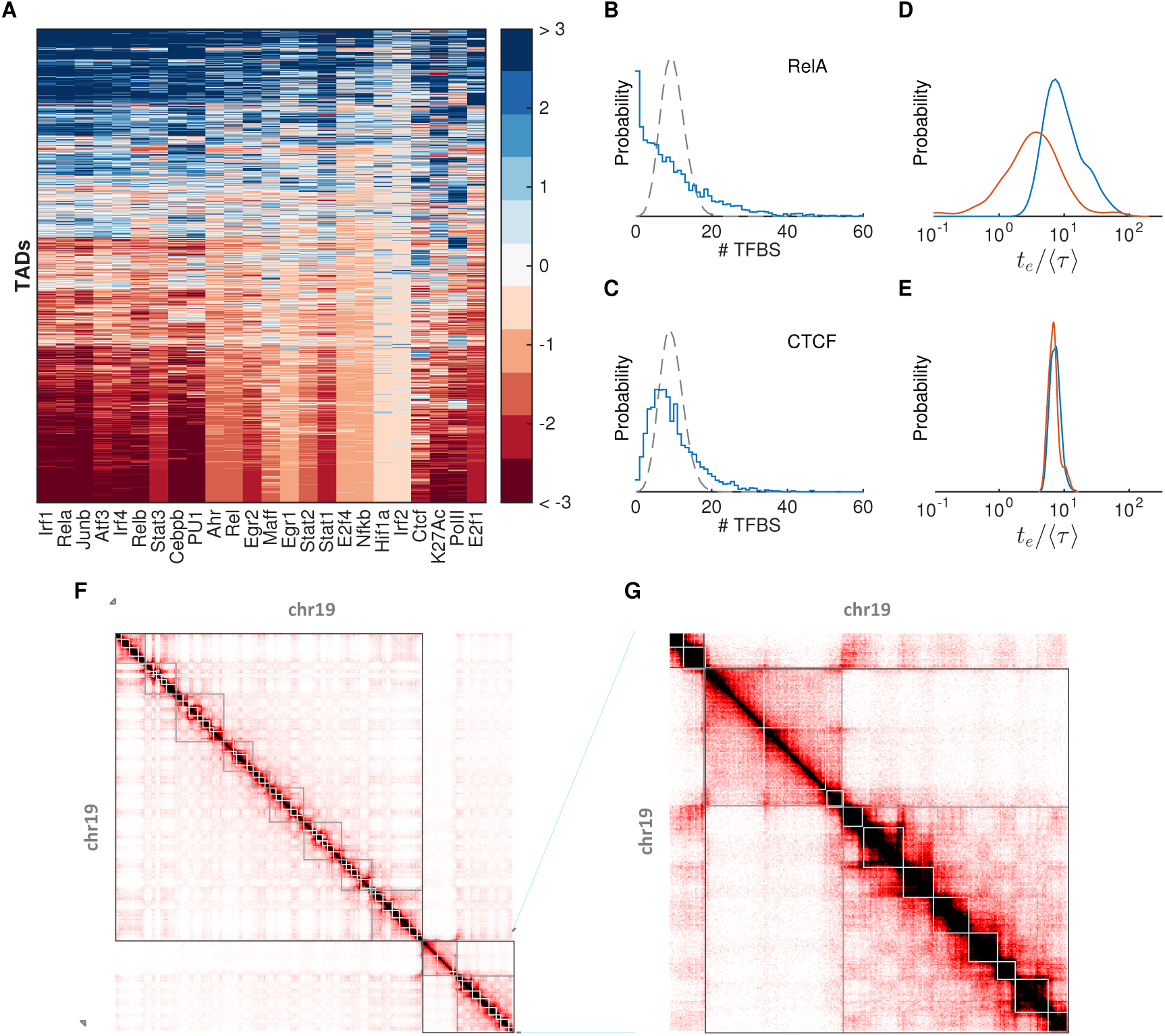
Asymmetric distribution of TFBSs across TADs traps TF diffusion. (A) Heat-map showing the overrepresentation (blue) or underrepresentation (red) of TFBSs across TADs. TFBSs cluster in few TADs. (B and C) Histograms showing the frequency of TADs with certain number of TFBSs (blue) compared with a random distribution (gray). Notably TADs with small number of sites or with large number of sites occur significantly more often than expected by a random model. (D) Distribution of normalized escaping times *t*_*e*_ /(τ) from TADs with one or zero sites (red) or with more than 15 sites of RelA. (E) Same as in D but with randomized residence times τ_i_. (F) Diffusive associated domains (DADs) for different time scales (white, grey and dark grey) obtained from stability analysis of a diffusion process on the chromatin network with non-specific DNA-interactions.

Conversely, the diffusion process can be used to study the chromatin structure at different scales. This idea has been already applied to identify hierarchical structures and communities in a wide variety of networks [16]. We defined diffusion associated domains (DADs) as chromatin structures from which a diffusive molecule is not likely to scape after certain time. In Fig. 4F and 4G we showed the obtained DADs at three different time scales. These results revealed in a clear an intuitive manner the hierarchical and fractal nature of the chromatin structure [17].

## Conclusions

We presented a stochastic model of TF diffusion that for the first time integrates high-resolution information on the 3D structure of chromatin and DNA-protein interaction. Notably, the multi-scale structure of our model allowed us to extend the description of the DNA-protein interaction by introducing easily genome-wide information on histone post-transcriptional modifications. Our model allowed us to uncover the effects of chromatin struc-ture on transcription factor occupancy profiles and target search times. Finally, we showed that TF-BSs clearly clustered preferentially in few TADs which leads to a higher local concentration of TFs as they are trapped in those TADs for longer times. We hypothesized this could be an optimal strategy to efficiently use the limited cellular transcrip-tional resources.

## Acknowledgments

NM conceived and designed the study and developed the model. NM and NA performed the research. NM wrote the paper.

## Author contribution

Nacho Molina thanks Samuel Zambrano, Felix Naef, Davide Mazza and Lukas Burger for useful commentsand suggestions.

